# Impaired oligodendrocyte maturation is an early feature in SCA3 disease pathogenesis

**DOI:** 10.1101/2021.11.18.468958

**Authors:** Kristen H. Schuster, Annie J. Zalon, Hongjiu Zhang, Danielle M. DiFranco, Nicholas R. Stec, Zaid Haque, Kate G. Blumenstein, Amanda M. Pierce, Yuanfang Guan, Henry L. Paulson, Hayley S. McLoughlin

## Abstract

Spinocerebellar ataxia type 3 (SCA3), the most common dominantly inherited ataxia, is a polyglutamine neurodegenerative disease for which there is no disease-modifying therapy. The polyglutamine-encoding CAG repeat expansion in the *ATXN3* gene results in expression of a mutant form of the ATXN3 protein, a deubiquitinase that causes selective neurodegeneration despite being widely expressed. The mechanisms driving neurodegeneration in SCA3 are unclear. Research to date, however, has focused almost exclusively on neurons. Here, using equal male and female age-matched transgenic mice expressing full-length human mutant *ATXN3,* we identified early and robust transcriptional changes in selectively vulnerable brain regions that implicate oligodendrocytes in disease pathogenesis. We mapped transcriptional changes across early, mid, and late stages of disease in two selectively vulnerable brain regions, the cerebellum and brainstem. The most significant disease-associated module through weighted gene co-expression network analysis revealed dysfunction in SCA3 oligodendrocyte maturation. These results reflect a toxic gain of function mechanism, as ATXN3 knockout mice do not exhibit any impairments in oligodendrocyte maturation. Genetic crosses to reporter mice revealed a marked reduction in mature oligodendrocytes in SCA3-disease vulnerable brain regions and ultrastructural microscopy confirmed abnormalities in axonal myelination. Further study of isolated oligodendrocyte precursor cells from SCA3 mice established that this impairment in oligodendrocyte maturation is a cell autonomous process. We conclude that SCA3 is not simply a disease of neurons and the search for therapeutic strategies and disease biomarkers will need to account for non-neuronal involvement in SCA3 pathogenesis.

**SIGNIFICANCE STATEMENT:** Despite advances in SCA3 disease understanding, much remains unknown about how the disease gene causes brain dysfunction ultimately leading to cell death. We completed a longitudinal transcriptomic analysis of vulnerable brain regions in SCA3 mice to define the earliest and most robust changes across disease progression. Through gene network analyses followed up with biochemical and histological studies in SCA3 mice, we provide evidence for severe dysfunction in oligodendrocyte maturation early in SCA3 pathogenesis. Our results advance understanding of SCA3 disease mechanisms, identify additional routes for therapeutic intervention, and may provide broader insight into polyglutamine diseases beyond SCA3.

## INTRODUCTION

Spinocerebellar ataxia type 3 (SCA3), also known as Machado Joseph Disease (MJD), belongs to the family of polyglutamine (polyQ) diseases that include Huntington’s disease (HD), SCA types 1, 2, 6, 7, and 17, Spinal and Bulbar Muscular Atrophy (SBMA), and Dentatorubral Pallidoluysian Atrophy (DRPLA)^1,2^. These dominantly inherited polyQ disorders are characterized by a trinucleotide CAG repeat expansion in the corresponding disease gene. In SCA3, the expanded CAG repeat is located within the *ATXN3* gene, encoding an expanded polyQ repeat in the ATXN3 protein. SCA3 is a progressive and ultimately fatal disease in which the expanded protein is toxic and precipitates a host of cellular dysfunctions^3^. While individual cellular dysfunctions are presumably harmful, it is likely the sum of dysregulated pathways that culminate in neurodegeneration. An effective preventive treatment currently does not exist for SCA3. Further understanding of the salient dysregulated pathways occurring early in SCA3 is needed in order for rational therapeutic strategies to be developed.

While mutant ATXN3 is ubiquitously expressed throughout all cell types, SCA3 is classified as a neurodegenerative disease due to its clinical presentation and neuropathological features. Symptoms of SCA3 include progressive ataxia, dystonia, rigidity, distal muscle atrophy, and dysarthria^4^. The most prominent and well-studied neuronal loss occurs in the substantia nigra, brainstem, cerebellum, and spinal cord^5–8^. Our recent transcriptomic analysis of 20-week-old pontine tissue from a series of SCA3 transgenic overexpression, SCA3 knock-in, and *ATXN3* knock-out (KO) mouse models supported the view that the repeat expansion in SCA3 acts in a dominant-toxic manner associated with expression of the aggregation-prone mutant protein^9^. In contrast, loss of ATXN3 function did not promote such transcriptional changes^9^, and *ATXN3* KO mice appear phenotypically normal^10^, suggesting that loss of ATXN3 function is not a significant pathogenic factor in SCA3.

ATXN3 normally localizes to both the cytoplasm and nucleus, but in disease mutant ATXN3 tends to aggregate in the nucleus^11^. This change in subcellular localization, which is also observed for other polyQ disease proteins, implies that nuclear events, such as gene expression, could be altered in disease. In fact, transcriptional profiling studies across different polyQ diseases do suggest that transcriptional dysregulation contributes to polyQ disease pathogenesis. Results from the HD, SCA1, SCA2, and SCA7 studies highlight the fact that the disease state transcriptome changes both spatially and temporally^12–15^. While informative, prior studies characterizing the SCA3 CNS transcriptome were limited to studying either a single mouse brain region at mid-stage disease^9^, across two timepoints in the cerebellum^16^, or exploring transcriptional changes across different brain regions only in late-stage disease^17^. Here, we evaluate the CNS transcriptome of an SCA3 mouse model at multiple time-points to better understand expression changes that correlate with disease progression in two vulnerable SCA3 brain regions.

We compared transgenic SCA3 mice expressing the full-length human mutant *ATXN3* gene to wildtype littermates to map the earliest SCA3 disease transcriptional changes. Through RNA sequencing, we longitudinally characterized the CNS transcriptome during early-, mid-, and late-stages of disease in two highly affected brain regions, the brainstem and cerebellum. Using weighted gene co-expression network analysis (WGCNA), we identified early and progressive disease-associated modules, with the most significant module enriched for oligodendrocyte transcripts. To dissect how this cell type contributes to SCA3 disease, we characterized the spatiotemporal dysregulation of oligodendrocytes in SCA3 mice and defined oligodendrocyte impairments to be both cell autonomous and occurring through a toxic gain of function mechanism. Our data supports the view that SCA3 is not simply a neuronal disease and the search for therapeutic strategies and disease biomarkers for this fatal disorder will need to account for nonneuronal involvement, especially oligodendrocytes early in disease progression.

## MATERIALS AND METHODS

### Mice

All animal procedures were approved by the University of Michigan Institutional Animal Care and Use Committee and conducted in accordance with the United States Public Health Service’s Policy on Humane Care and Use of Laboratory Animals. Genotyping was performed using tail biopsy DNA isolated prior to weaning and confirmed postmortem, as previously described^18^. To determine the *ATXN3* CAG-trinucleotide repeat lengths in SCA3 homozygous Q84 mice^19^, DNA were analyzed by gene fragmentation analysis (Laragen Inc., Culver City) using *ATXN3* primers (5’-ACAGCAGCAAAAGCAGCAA-3’ and 5’-CCAAGTGCTCCTGAACTGGT-3’). CAG repeat length was calculated as (peak amplicon fragment size – 66) / 3, with the mean +/− SD repeat size of 75.625 +/− 2.24 for brainstem samples and 75.53 +/− 2.29 for cerebellar samples. Genotyping for *Mobp-*eGFP GENSET line^20^ (MGI:4847238) was completed using *Mobp* primers (5’-GGTTCCTCCCTCACATGCTGTTT-3’ and 5’-TAGCGGCTGAAGCACTGCA-3’). Genotyping for the *ATXN3* KO line^21^ was completed using ATXN3KO primers (For 5’-GAGGGAAGTCGTCATAAGAGT-3’, ATXN3KO Rev 5’-TGGGCTACAAGAAATCCTGTC-3’, and ATXN3KO LTRa 5’-AAATGGCGTTACTTAAGCTAG-3’). Animals were sex-, littermate- and age-matched for each study. For transcriptional profiling and histological analyses, 4-6 animals (equal number of males and female) per genotype were euthanized either at 3, 4, 8, 16, or 65 weeks of age and left-hemisphere tissue was macro-dissected into brainstem and cerebellum for RNA and protein, as previously described^18^. Right hemisphere tissue was post-fixed in 4% paraformaldehyde for histological preparations, as previously described^18^. Experimenters were blinded to genotypes during all analyses.

### RNA Isolation and Sequencing

RNA was extracted from PBS-perfused, flash-frozen brainstem and cerebellar samples from the left hemisphere. Using the Next Advance Bullet Blender, samples were homogenized in radioimmunoprecipitation assay (RIPA, by Sigma) buffer (750 μL for brainstem and 500 μL for cerebellum) with added proteinase inhibitor (BioRad). A 250 μL aliquot of this lysate was reserved for protein analysis and the remaining sample was used for RNA extraction. RNA extraction was carried out using the QIAshredder (Qiagen), RNeasy Mini Kit (Qiagen), and DNAse I kit (Qiagen), then eluted in RNase-free water, according to manufacturer’s instructions. RNA samples were assessed by nanodrop to determine concentration and sample purity. 1 μg of total RNA was submitted to the University of Michigan Sequencing Core for Illumina HiSeq library preparation, quality control, and RNA sequencing. We chose wildtype and Q84/Q84 RNA samples for RNA sequencing based on RIN number (>7 RIN) and, if applicable, CAG repeat expansion size (> average 72Q) as determined by Laragen, Inc. (Culver City, CA, USA).

### RNA-seq Expression Analyses

Purified RNA samples were submitted to the University of Michigan Bioinformatics Core for library generation (Illumina TruSeq), QC analysis, and sequencing on Illumina HiSeq 4000 (75×75 paired-end, 4 lanes). The RNA-seq data were pseudo-aligned to the ENSEMBL 94 mouse reference cDNA sequences (Oct 2018, GRCm38p6) using Kallisto 0.45. The transcript quantification results from Kallisto were then fed to Sleuth 0.30 for differential expression analysis, aggregated by their genes.

From samples collected from each tissue type (brainstem or cerebellum) at each time point (8, 16, or 65 weeks), genes with expression levels less than 5 transcripts per million (TPM) in more than 25% of the analyzed samples were discarded from the differential expression analysis. For each group, Sleuth evaluated the gene-wise statistical significance of the hypothesis that a gene is differentially expressed in wildtype samples from transgenic samples. Through Benjamini-Hochberg correction, genes with q-values less than 0.05 were collected as the differentially expressed genes. The WGCNA R package version 1.66 was used to construct a gene co-expression network with the default being a soft threshold power [beta] of 6^22^. Eighteen modules were detected, including three that were significantly associated with SCA3 (wildtype versus Q84/Q84 mice, t-test, Bonferroni corrected p value < 1e-5). All differentially expressed genes identified in the Turquoise WGCNA modules were used to identify significant upstream regulators and Tox lists using Ingenuity Pathway Analysis (IPA)^23^.

### Quantitative Real-Time PCR

Validation of RNA-seq results by quantitative real-time PCR (qPCR) analysis was performed through reverse transcription on 0.5-1 μg of total RNA using the iScript cDNA synthesis kit according to the manufacturer’s instructions (Bio-Rad, Hercules, CA). The cDNA was diluted 1:20 in nuclease-free water. iQ SYBR green qPCR was performed on the diluted cDNA following the manufacturer’s protocol (Bio-Rad, Hercules, CA) using previously described primers^9,18^ or the following primers: human *ATXN3* (Forward primer: CCTCAATTGCACATCAGCTGGAT; Reverse primer: AACGTGCGATAATCTTCACTAGTAACTC; Probe Seq: CTGCCATTCTCATCCTC), mouse *Atxn3* (Thermofisher; Cat # Mm01336273), *Beta Actin* (Thermofisher; Cat # Mm02619580), *Smoc1* (Thermofisher; Cat #Mm00491564), *Dusp15* (Thermofisher; Cat #Mm00521352), *Aspa* (For 5’-TTCCACTGGGTGGAGACTGT-3’; Rev 5’- TTCCACTGGGTGGAGACTGT-3’), *Ugt8a* (Thermofisher; Cat #Mm00495930), *Plp1* (Thermofisher; Cat #Mm01297210), *Mobp* (Thermofisher; Cat #Mm02745649), *Mal* (Thermofisher; Cat #Mm01339780). Gene expression fold change (RQ) was calculated relative to Q84/Q84 samples for each timepoint and normalized to beta actin, where RQ = 2^−(ddCT)^.

### Immunohistochemistry

Perfused brains were post-fixed overnight in 4% paraformaldehyde (PFA) before switching to 30% sucrose in 1x phosphate buffered saline (PBS). Brains were cut via microtome into a series of 40μm sagittal sections free-floating into cryostorage as previously described^18^. For immunofluorescence studies, sections were washed 3 times with 1x PBS then incubated for 30 minutes at 80°C in 0.01M sodium citrate buffer, pH 8.5, for antigen retrieval. Sections were again washed with PBS and blocked for one hour at room temperature (RT) in M.O.M. Mouse Ig Blocking Reagent (Vector Laboratories, BMK-2202) diluted in PBS with 5% normal goat serum (NGS) according to manufacturer instructions. After blocking, sections were washed twice in PBS, incubated for 15 minutes in M.O.M. Diluent in PBS according to manufacturer instructions, and incubated overnight in primary antibody at 4°C. Primary antibodies assessed include mouse anti-ATXN3 (1H9) (1:500; MAB5360; Millipore) and rabbit anti-Olig2 (1:500; P21954; ThermoFisher). Sections were washed 3 times with PBS, followed by incubation with Alexa Flour 568 goat anti-mouse IgG (1:1000; A11011; Invitrogen) and Alexa Fluor 647 goat anti-rabbit (1:1000; A21245; Invitrogen) secondary antibodies. All sections were stained with DAPI (Sigma) for 15 min at RT and mounted with Prolong Gold Antifade Reagent (Invitrogen). Imaging was performed using a Nikon-A1 confocal microscope in the basilar pontine nuclei (denoted as pons), deep cerebellar nuclei (DCN), corpus callosum (CC), or corticospinal tract (CST). Images were analyzed using CellProfiler software ^24^.

### Western blot

Protein lysates from macrodissected brainstem and cerebellar tissue were processed as previously described^18^ and stored at −80°C. Thirty micrograms of total mouse brain protein lysate were resolved in 4 to 20% gradient sodium dodecyl sulfate–polyacrylamide electrophoresis gels and transferred to 0.45 μm nitrocellulose membranes. Membranes were incubated overnight at 4°C with MOBP (Rabbit; 1:100; ab91405; Abcam) and GAPDH (mouse; 1:5000; MAB374; Sigma) primary antibodies. Molecular weight bands were visualized by incubation with peroxidase-conjugated anti-mouse or anti-rabbit secondary antibody (1:5000; Jackson Immuno Research Laboratories) followed by treatment with the ECL-plus reagent (Western Lighting; PerkinElmer) and exposure on G:Box Chemi XRQ system (Syngene). Band intensities were quantified using ImageJ analysis software (NIH).

### Transmission Electron Microscopy

Mice (1 male and 2 females per genotype) were transcardially perfused first with 1x PBS, then with 2.5% glutaraldehyde and 4% PFA in 1x PBS. Brains were removed and postfixed overnight at 4°C in the same buffer before being transferred to 0.1M phosphate buffer, pH 7.4 for long term storage at 4°C. Processing of TEM tissue was done by University of Michigan’s Microscopy Core according to their protocol. Briefly, post-fixed tissued was *en bloc* stained in 2% aqueous uranyl acetate, washed and dehydrated, then infiltrated with a graded resin series before polymerizing. Polymerized resin blocks were ultra-thin sectioned at 70nm using a DiATOME Ultra 45 diamond knife (DiATOME, Hatfield, PA) and a Leica EM UC7 ultramicrotome (Leica Microsystems Inc., Buffalo Grove, IL). Ultra-thin sections were imaged on a JEOL JEM-1400 plus transmission electron microscope (JEOL USA Inc., Peabody, MA) at an accelerating voltage of 60 kV, using either an AMT 4-megapixel XR401 or a 12-megapixel Nanosprint CMOS camera (Advanced Microscopy Techniques, Woburn, MA). For g-ratio, axon caliber, and cumulative fraction analysis of myelinated axons, MyelTracer software was used^25^.

### OPC Isolation and Culture

Oligodendrocyte precursor cells (OPCs) were isolated from 5–7-day old YACQ84 transgenic mice using magnetic cell sorting (MACS, Miltenyi Biotec, Bergisch Gladbach, Germany). Whole brains were dissected and homogenized using the Neural Tissue Dissociation Kit P (Miltenyi, #130-092-628) and gentleMACS dissociator (Miltenyi, #130-093-235). OPCs were magnetically labeled with microbeads conjugated to monoclonal PDGFRα (Miltenyi, #130-101-502) and isolated using LS columns (Miltenyi, #130-042-401). OPCs were plated in 8-well slides at 5×10^3^ per mm^2^ in OPC media (+FGF2, +PDGFAA). The following day, OPCs were either fixed with 4% PFA for 20 minutes at room temperature (DIV0) or switched to differentiation media (OPC media, −FGF2, −PDGFAA, +T3) for collection at DIV3.

### Immunocytochemistry

After fixation, OPCs were washed once with 1x PBS, blocked for 30 minutes at RT with 5% NGS in PBS, and permeabilized with 0.5% Triton X-100 in blocking buffer for 10 minutes. Cells were then incubated in primary antibodies overnight at 4°C. Primary antibodies used include rabbit anti-SMOC1 (1:100, PIPA531392, ThermoFisher), rat anti-MBP (1:200, ab7349, Abcam), and mouse anti-ATXN3 (1H9) (1:500, MAB5360; Millipore). Following primary incubation, OPCs were washed twice with PBS and incubated with corresponding AlexaFluor488, 568, or 647 secondary antibodies (1:1000, Invitrogen) diluted in blocking buffer for 1 hour at RT. Cells were again washed with PBS, stained with DAPI (Sigma) for 15 minutes at RT, and mounted with Prolong Gold Antifade Reagent (Invitrogen). Imaging was performed using a Nikon-A1 confocal microscope. Four images were randomly taken in each quadrant of a well, with 2 wells per mouse, and 3-4 mice per genotype. Images were analyzed using CellProfiler software^24^.

### Statistics

All statistical analyses were performed using Prism (8.0; GraphPad Software, La Jolla, CA). All statistical significance was tested using either a student’s t-test or a one-way ANOVA with a post hoc Tukey’s multiple comparisons test. Variability about the mean is expressed as mean ± standard error of the mean (SEM). All tests set the level of significance at p<0.05.

## RESULTS

### SCA3 mice exhibit different temporal and regional transcriptional profiles relative to wildtype littermates

Many *in vitro* and *in vivo* model systems of SCA3 have been employed to study the molecular consequences of mutant (polyQ-expanded) ATXN3 expression. For this study, we used the well-characterized YACMJD84.2Q-C57BL/6 (Q84) transgenic mouse line, first described in 2002^19^, because it expresses the full-length human mutant *ATXN3* gene ubiquitously via its endogenous promoter. Hemizygous Q84 mice harbor two copies of the transgene, initially described with an expanded CAG repeat length of 84, which can be bred to homozygosity to elicit a stronger phenotype. The presence of the full-length *ATXN3* gene in Q84 mice ensures that all human isoforms of *ATXN3* are expressed^19^.

The Q84 mouse recapitulates many disease-relevant phenotypes making it ideal for molecular studies of SCA3 pathogenesis. Homozygous Q84 mice develop motor and behavioral phenotypes similar to deficits described in SCA3 patients including abnormal gait, coordination deficits, muscle weakness, reduced weight gain, and premature death. They also share critical neuropathological features of disease including intranuclear accumulation of ATXN3 in neurons and the formation of ATXN3 aggregates widely in the brain^19,26,27^.

To define transcriptional changes occurring over time, we harvested tissue from Q84/Q84 mice and wildtype (WT) littermates at 8, 16, and 65 weeks. These ages correspond to early-, mid-, and late-stage disease as defined by ATXN3 nuclear accumulation pathology and motor assessment, based on published studies^26,27^. We extracted RNA from two highly affected brain regions in SCA3 disease, the brainstem and cerebellum. Samples with RNA integrity number (RIN) greater than 7 were submitted for paired end RNA-sequencing (RNA-seq). The average number of read counts per brainstem sample was 48.8 million (SD ± 7.3 million), with an average of 84% of sequencing reads mapping to exons of known genes. The average number of read counts per cerebellum sample was 31.1 million (SD ± 3.2 million), with an average of 83% of sequencing reads mapping to exons of known genes.

Consistent with a recent end-stage transcriptomic study of this SCA3 mouse model^17^, our RNA-seq data revealed altered gene expression in both the brainstem and cerebellum, with more robust changes occurring in the brainstem at all tested ages (**Fig. 1**). Bioinformatic analysis of the differentially expressed (DE) genes shared across all three timepoints identified 43 genes in the brainstem and 6 in the cerebellum (**Fig. 1B**). Not unexpectedly, the total number of DE genes markedly increased with age and disease progression. Roughly 25-30% of DE genes were upregulated and 70-75% downregulated in both brain regions.

**Figure 1.**
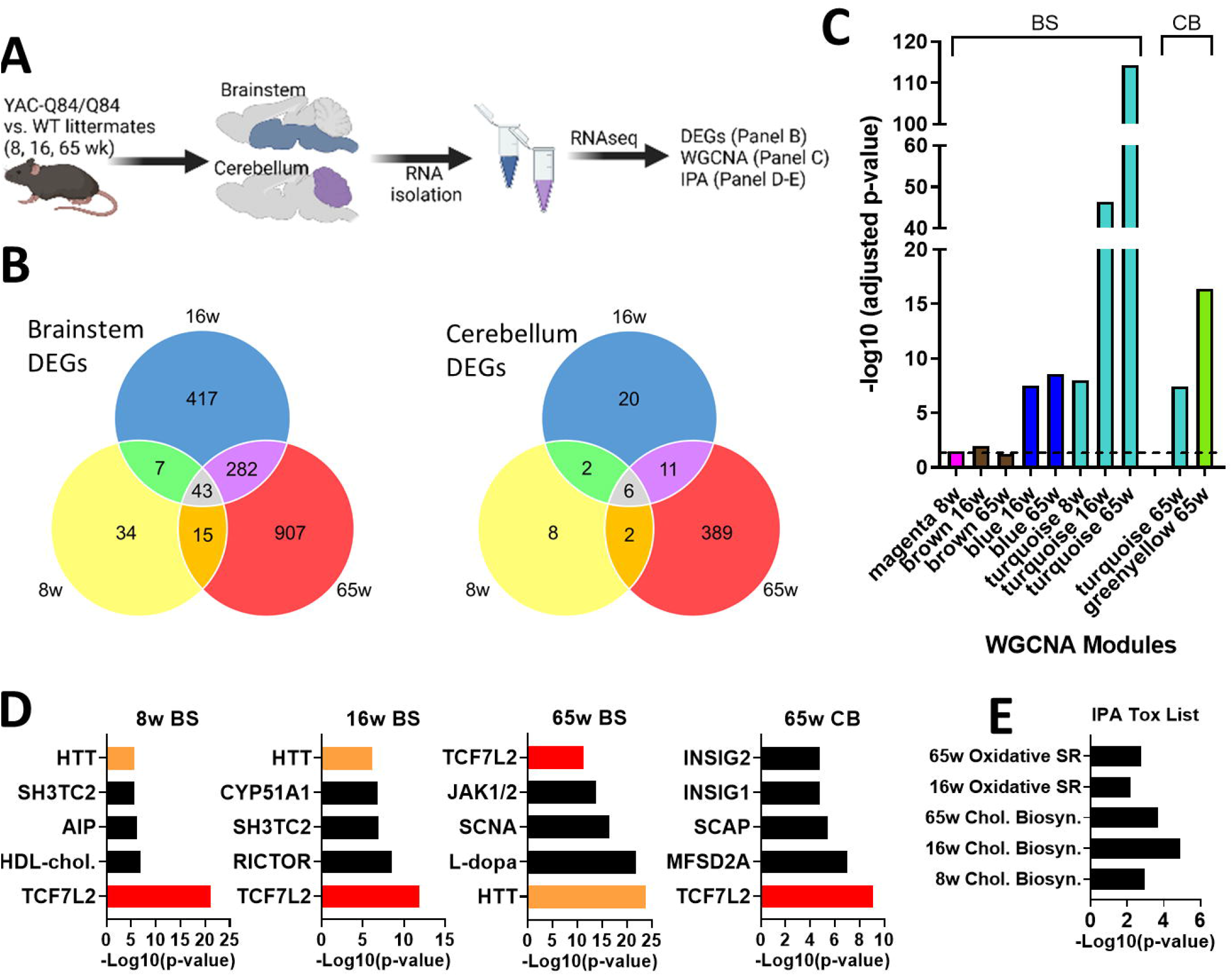
Spatiotemporal gene co-expression networks implicated in SCA3 mice. (A) Schematic pipeline of SCA3 RNA-seq and analysis of mouse brainstem and cerebellum samples (n=5-6 mice/timepoint/genotype). (B) Number of differentially expressed genes (DEGs) between WT and Q84/Q84 mice at 8, 16, and 65 weeks of age in the brainstem and cerebellum. (C) Identification of significant brainstem (BS) and cerebellum (CB) weighted gene co-expression network analysis (WGCNA) modules in SCA3 mice relative to age-matched controls. (D) Lists of top five upstream regulators identified by Ingenuity Pathway Analysis (IPA) include TCF7L2 (red) and HTT (orange) in brainstem at all timepoints and in cerebellum at 65 weeks. (E) Significant IPA Tox list results across brainstem timepoints (Oxidated SR=Oxidative Stress Response; Chol. Biosyn=Cholesterol Biosynthesis).

### A disease associated WGCNA module is enriched for oligodendrocyte genes

Using WGCNA^22^, we identified gene sets whose expression significantly correlate in at least one of the age groups and at least one of the tested brain regions. Among the co-expression modules, Blue, Turquoise, and Greenyellow modules were highly significant in SCA3 disease tissues relative to WT littermates (**Fig. 1C**). The Turquoise module was detected at all ages in the brainstem, increasing in significance as disease progressed. This module also proved to be significant in the cerebellum at late-stage disease.

To evaluate the biological relevance of the Turquoise module, we performed Ingenuity Pathway Analysis (IPA)^23^. Within this module, a key regulator of oligodendrocyte differentiation^28–32^, transcription factor 7-like 2 (TCF7L2), was among the top five IPA-defined upstream regulators at all tested ages (**Fig. 1D**). Until recently, TCF7L2 had been described exclusively as a Wnt signaling effector for myelin formation^30^. In 2016, Zhao et al. ascribed another function to TCF7L2: a dual-regulatory switch in oligodendrocyte maturation^29^. TCF7L2 can either interact with beta-catenin to induce Wnt transcription, thereby inhibiting differentiation, or can form a complex with Kaiso/Zbtb33 and Sox10 to induce oligodendrocyte differentiation^29^. Additionally, a 2017 study in which *Tcf7l2* was conditionally deleted in mice showed TCF7L2 to be essential for promoting oligodendrocyte differentiation and re-myelination. Because TCF7L2 is preferentially expressed in newly forming oligodendrocytes, deletion of the gene had no effect on oligodendrocyte precursor cells (OPCs)^33^.

In addition to TCF7L2, the HD gene Huntingtin (HTT) was identified in the brainstem as a significant upstream regulator in the Turquoise module at all ages (**Fig. 1D**). While little is known about mutant ATXN3 behavior in oligodendrocytes, recent studies in the HD field describe delayed oligodendrocyte differentiation that can be resolved by reducing HTT expression in the oligodendroglial lineage^34,35^, implying that oligodendrocyte dysregulation is cell autonomous in HD. Consistent with these findings, our IPA Tox analysis also suggested impaired cholesterol biosynthesis pathways, which are essential for myelin growth^36^ (**Fig. 1E**).

To explore the relationship between oligodendrocyte dysregulation and disease progression, we identified 34 Turquoise module genes that were consistently differentially expressed across all tested ages in SCA3 brainstem tissue (**Fig. 2A**). When querying gene expression patterns using an existing RNA sequencing database of immune-panned brain cells^37^, we found that more than 60% of the overlapping genes were highly expressed in oligodendrocytes and expressed at low levels in neurons, astrocytes, and microglia (**Fig. 2A**).

**Figure 2.**
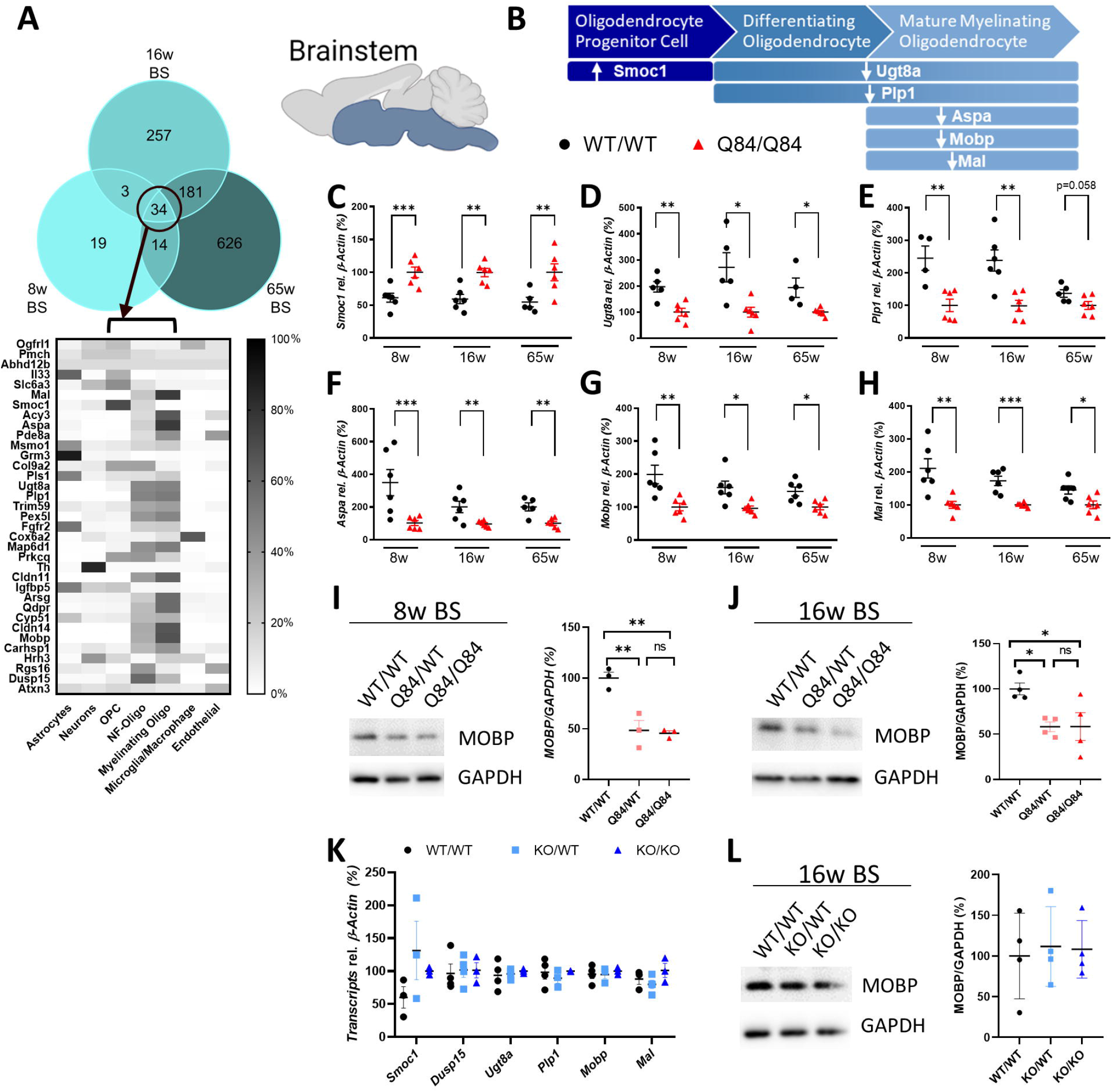
Dysregulation of oligodendrocyte genes in SCA3 brainstem tissue. (A) Reported relative cell-type expression of the 34 Turquoise module genes that were consistently differentially expressed in SCA3 mice; cell type expression is based on RNA-seq of immunopanned mouse brain cells^34^. OPC= oligodendrocyte precursor cells, NF-Oligo= newly forming-oligodendrocytes. The turquoise module from RNA sequencing analysis is enriched for oligodendrocyte lineage genes, which are dysregulated in the brainstem of SCA3 mice. (B) Illustrative representation of oligodendrocyte development and directional gene change in Q84/Q84 mice relative to wildtype littermates. (C-H) QPCR confirmation of brainstem DEGs pertaining to individual oligodendrocyte developmental stages between Q84/Q84 and WT mice at 8,16, and 65 weeks of age. Validated genes include *Smoc1, Ugt8a, Plp1, Aspa, Mobp,* and *Mal.* Data (mean ± SEM) are reported relative to Q84/Q84 samples (n= 4-6 per condition). Unpaired parametric t-tests, assuming equal SD, were performed (*p < 0.05, **p <0.01, ***p <0.001). (I-J) Representative western blot image and quantification of a mature oligodendrocyte marker (MOBP) in 8- and 16-week-old brainstem tissue (n=3-4 mice per genotype) analyzed by one-way ANOVA with Tukey’s multiple comparisons test (*p < 0.05, **p <0.01, ns=not significant). (K) QPCR analysis of oligodendrocyte development genes in the brainstem show no differences in ATXN3 KO tissue in all transcripts assessed by one-way ANOVA with Tukey’s multiple comparisons test. (L) Similarly, no changes were found in MOBP expression levels by western blot analysis in ATXN3 KO brainstem tissue by one-way ANOVA with Tukey’s multiple comparisons test.

### Oligodendrocyte maturation is impaired via a toxic gain of function mechanism

Oligodendrocyte maturation can be divided into three stages: OPCs, differentiating or newly formed oligodendrocytes, and mature myelinating oligodendrocytes (**Fig. 2B**). Among the 34 consistently DE genes in SCA3 mouse brainstem, we chose six genes known to exhibit high oligodendrocyte expression for analysis by qPCR: *Smoc1, Ugt8a, Plp1, Aspa, Mobp,* and *Mal.* In SCA3 mice, there was upregulation of *Smoc1*, which is known to be highly expressed in OPCs, whereas genes known to be expressed in differentiating oligodendrocytes (e.g., *Ugt8a and Plp1*) and mature oligodendrocytes (e.g., *Aspa, Mobp, and Mal*) were downregulated (**Fig. 2C-H**). Furthermore, western blot quantification of the mature oligodendrocyte marker (MOBP) in 8- and 16-week-old brainstem tissue correlated with transcriptional changes: approximately a 50% reduction of MOBP protein in SCA3 mice compared to wildtype littermates (**Fig. 2I-J**). In comparison, cerebellar transcriptional expression of oligodendrocyte genes showed similar trends, however differences across timepoints were less consistent and significant (**Fig. 3A-E**). Protein expression of MOBP in cerebellar tissue shows a dose-dependent reduction in 8- and 16-week disease samples compared to wildtype littermates (**Fig. 3F-G**). These transcriptional and biochemical analyses implicate a role for mutant ATXN3 in regulating oligodendrocyte maturation with more severe dysregulation in brainstem tissue relative to cerebellar tissue.

**Figure 3.**
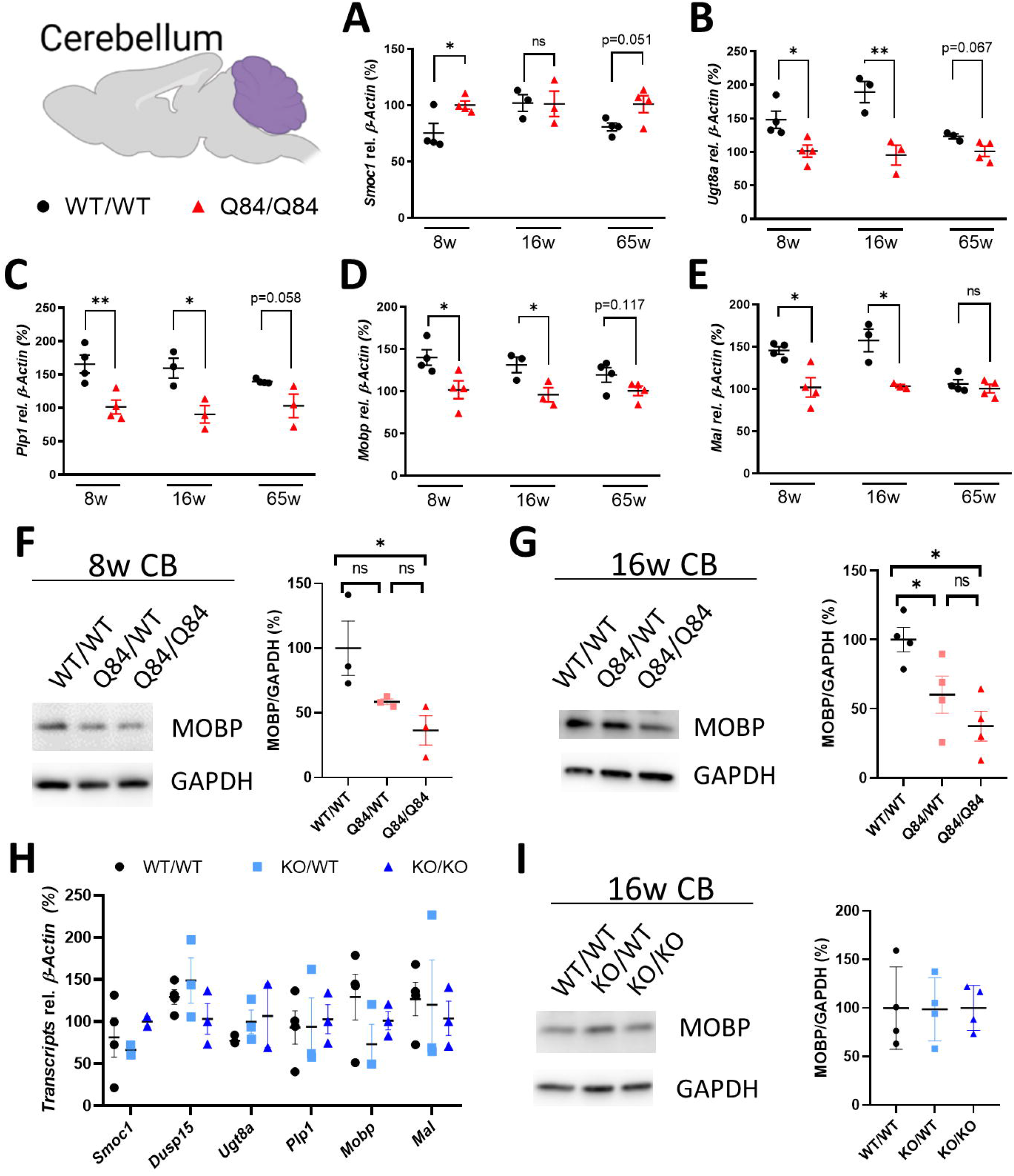
Oligodendrocyte maturation genes also dysregulated in SCA3 mouse cerebellar tissue. (A-E) Validation by qPCR of gene expression changes over time in SCA3 cerebellum corresponding to oligodendrocyte genes across developmental stages. Validated genes include *Smoc1, Ugt8a, Plp1, Mobp,* and *Mal.* Data (mean ± SEM) are normalized to Q84/Q84 samples (n= 4-6 per condition). Unpaired parametric t-tests, assuming equal SD, were performed (*p < 0.05, **p <0.01, ns=not significant). (F-G) Representative western blot image and quantification of MOBP, a mature oligodendrocyte marker, in cerebellar tissue at 8 and 16 weeks of age (n=3-4 mice per genotype) analyzed by one-way ANOVA with Tukey’s multiple comparisons test (*p < 0.05, ns=not significant). (K) QPCR and (L) western blot analyses of oligodendrocyte maturation markers at 16 weeks show no changes in the cerebellum of ATXN3 KO mice by one-way ANOVA with Tukey’s multiple comparisons test.

To address if loss of ATXN3 function also contributes to oligodendrocyte maturation impairment, we collected brainstem and cerebellar tissue from 16-week-old heterozygote (KO/WT) and homozygote (KO/KO) *ATXN3* knockout mice, and wildtype littermates. QPCR analysis of oligodendrocyte development genes show no change between wildtype, KO/WT, or KO/KO in all transcripts assessed (**Fig. 2K and 3H**). Similarly, no changes were found in MOBP expression levels by western blot analysis in either brain region (**Fig. 2L and 3I**). These results demonstrate that loss of ATXN3 function does not contribute to impaired oligodendrocyte maturation early in SCA3 disease.

### Oligodendrocyte maturation in SCA3 mice is impaired in selectively vulnerable brain regions

Guided by the oligodendrocyte transcriptional and biochemical results, we tested whether the observed changes in oligodendrocyte gene expression were due to a reduction of oligodendrocytes in SCA3 disease tissue. To assess the number of mature oligodendrocytes in the SCA3 brain, we crossed SCA3 mice with the BAC transgenic *Mobp*-eGFP reporter mouse^20^, which allows for visualization of mature oligodendrocytes. Brain sections harvested from hemizygous and homozygous SCA3;*Mobp*-eGFP+ and wildtype;*Mobp*-eGFP+ littermate mice at 16 weeks were immunofluorescently labeled with anti-Olig2 and anti-ATXN3 antibodies, then co-stained with DAPI. Confocal images from the body of the pons, DCN, and corpus callosum (CC) were captured. In both vulnerable brain regions in SCA3 mice (pons and DCN), Olig2+ cell counts were not significantly different from wildtype littermates (**Fig. 4A-D**). Both Olig2 and ATXN3 are expressed in all developmental stages of oligodendrocyte maturation^37^, therefore this result implies there is no overt loss of oligodendrocyte lineage cells in a symptomatic SCA3 mouse. Our previous work in this mouse model also supports this result, as we found no increase in the cell death markers BAX and BCL-2 in analogous 12-week-old SCA3 brainstem tissue^27^. Interestingly, Olig2+ cell counts in SCA3 mice in a less-vulnerable brain region, the CC, were significantly increased relative to wildtype littermates (**Fig. 4E-F**). Because our data indicates the changes in oligodendrocyte maturation markers in disease mice are not due to a loss of oligodendrocytes, we questioned the capacity of these cells to reach a mature, myelinating state in vulnerable regions.

**Figure 4.**
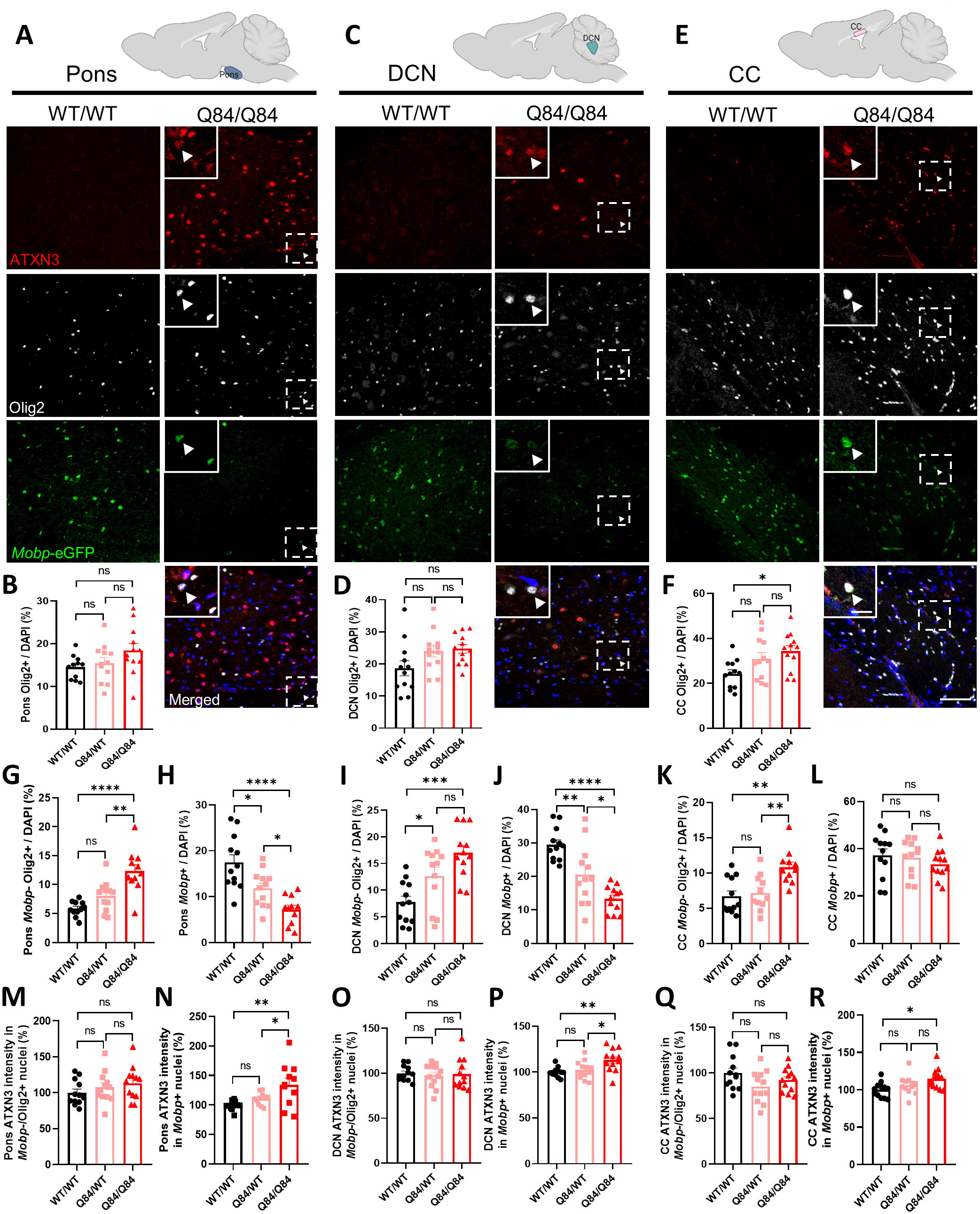
Reduction of mature oligodendrocytes occurs only in SCA3 vulnerable brain regions. To visualize mature oligodendrocytes in brain tissue, SCA3 mice were crossed to *Mobp*-eGFP reporter mice. (A, C, E) Representative sagittal immunofluorescent images of ATXN3 (red), Olig2 (white), and *Mobp*-eGFP (green) expression in 16-week-old Q84/Q84 mice and wildtype littermates in the body of the pons, deep cerebellar nuclei (DCN), and corpus callosum (CC). Scale bar, 50 μm. Inset scale bar, 12.5 μm. White arrowheads define cells with colocalization of nuclear ATXN3, Olig2, and MOBP. (B, D, F) No changes in total oligodendrocyte lineage cell counts (Olig2+ cells) were found in the pons or DCN, though a significant increase in Q84/Q84 mice compared to wildtype was noted in the CC. (G, I, K) Immature oligodendrocytes (Mobp-/Olig2+) were significantly increased in all three brain regions. (H, J, L) Mature oligodendrocytes (Mobp+) in the pons and DCN were significantly decreased, whereas no differences in mature oligodendrocyte cell counts were found in the CC. (M-R) ATXN3 accumulated in the nucleus of mature oligodendrocytes, but not in immature oligodendrocytes in all regions assessed. Cell counts normalized to total DAPI-stained nuclei per field; ATXN3 nuclear accumulation normalized to WT/WT images (n=3 images per mouse, n= 4 mice per genotype). Data reported as mean ± SEM. One-way ANOVA with Tukey’s multiple comparisons test was performed (*p < 0.05, **p <0.01, ***p <0.001, ****p<0.0001, ns=not significant).

To assess the number of non-myelinating (immature) versus myelinating (mature) oligodendrocytes in the SCA3 brain, we counted Olig2+ and *Mobp*-eGFP-positive cells in the pons, DCN, and CC. In all regions assessed, non-myelinating immature oligodendrocytes (Mobp-/Olig2+) were increased in SCA3 mice relative to wildtype littermates (**Fig 4G, I, K**). Strikingly, only in SCA3 vulnerable brain regions, the DCN and pons, were *Mobp-*eGFP-positive cells significantly decreased in Q84 mice compared to wildtype littermates (**Fig. 4H, J**). By contrast, no difference in *Mobp-*eGFP-positive cells counts were found in the CC of SCA3 versus wildtype mice (**Fig. 4L**). These results imply there is a tissue specific reduction of mature oligodendrocytes in SCA3 vulnerable brain regions.

An early and critical step in SCA3 pathogenesis is the redistribution of polyQ-expanded ATXN3 from the cytoplasm into neuronal nuclei^1,38^. While ATXN3 is known to be ubiquitously expressed across all CNS cell types, we found evidence of intranuclear ATXN3 accumulation in oligodendrocytes in 16-week-old SCA3 mice (**Fig. 4A, C, E inset;** white arrowheads). Furthermore, we quantified nuclear ATXN3 accumulation in non-myelinating oligodendrocytes (Mobp-/Olig2+) and myelinating oligodendrocyte (Mobp+) and across all regions found no difference in nuclear ATXN3 expression in non-myelinating oligodendrocytes but did observe dose-dependent increases of nuclear ATXN3 expressing in myelinating oligodendrocytes (**Fig. 4M-R**). Interestingly, this increased nuclear ATXN3 in myelinating oligodendrocytes was more significant in vulnerable brain regions (pons and DCN) than in nonvulnerable regions (CC).

### Reduction in mature myelinating oligodendrocytes cell counts correspond to ultrastructural abnormalities in the corticospinal tract

To understand the ultrastructural myelin changes occurring in our SCA3 mice, we sought out vulnerable SCA3 white matter tracts. A recent publication by Inada and colleagues defined through diffusion tensor imaging that microstructural damage to the corticospinal tracts (CST) is correlated with Scale for Assessment and Rating of Ataxia (SARA) in a large cohort of SCA3 patients^39^. Using the *Mobp-*eGFP reporter mouse crossed to the SCA3 mice, we explored oligodendrocyte maturation in the CST (**Fig 5A**) and again found no change in total Olig2+ cell counts between WT and SCA3 mice (**Fig 5B**). However, when quantified, we identified increased immature oligodendrocyte cell counts (**Fig. 5C**) and decreased mature oligodendrocyte cell counts (**Fig. 5D**) in the CST of SCA3 disease mice relative to wildtype littermates in a dose-dependent manner. Similar to the DCN and pons tissue, only mature myelinating oligodendrocytes, and not immature oligodendrocytes within the CST were found to have increased nuclear ATXN3 accumulation (**Fig. 5E-5F**). This result led us to collect tissues for ultrastructural analysis of myelin changes in the CST of SCA3 mice relative to wildtype littermates.

**Figure 5.**
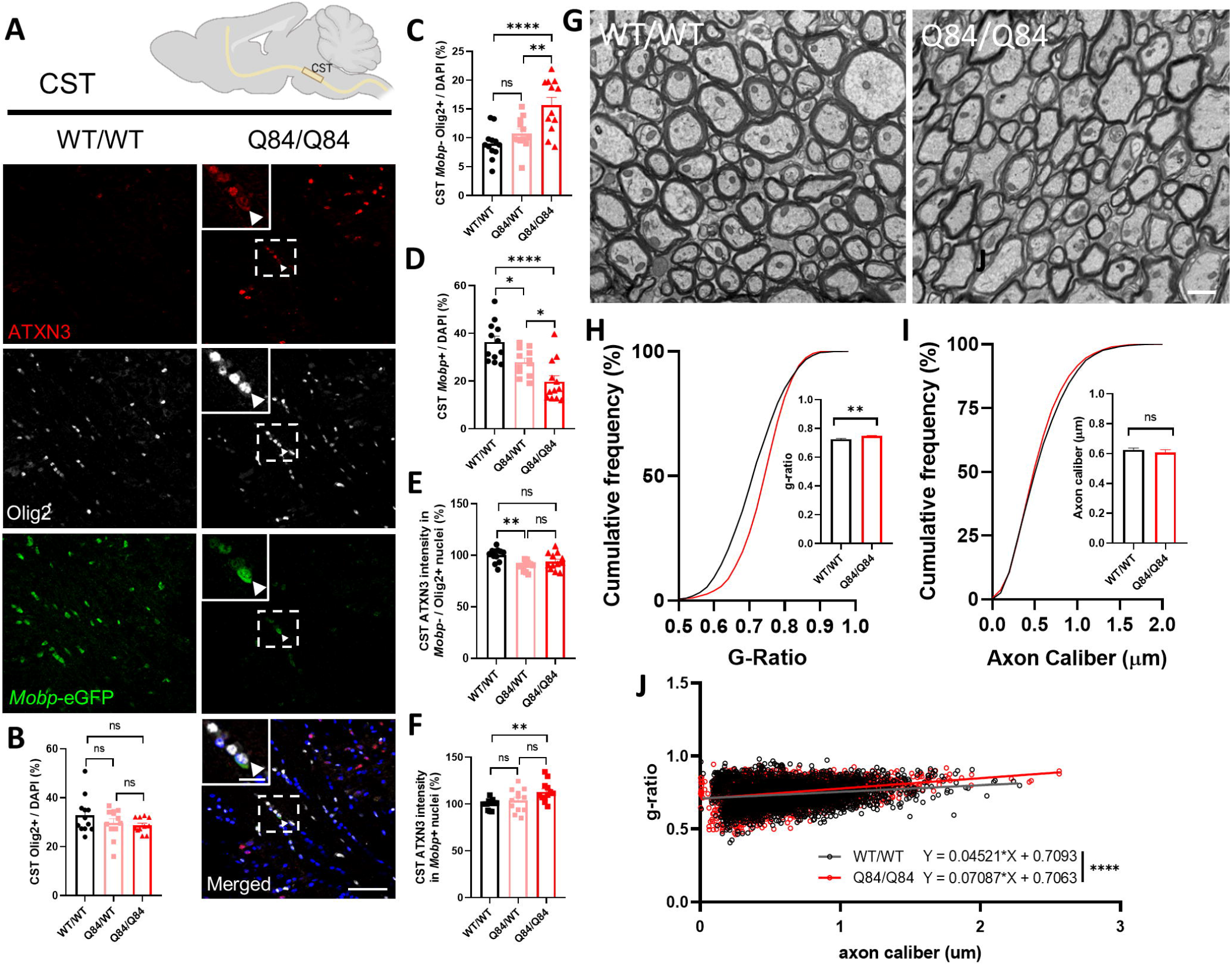
Reduction in *Mobp*-eGFP+ oligodendrocytes corresponds to ultrastructural abnormalities in the corticospinal tract. (A) Representative sagittal immunofluorescent images of ATXN3 (red), Olig2 (white), and *Mobp*-eGFP (green) expression in 16-week-old Q84/Q84 mice and wildtype littermates in the corticospinal tract (CST). Scale bar, 50 μm. Inset scale bar, 12.5 μm. White arrowheads define cells with colocalization of nuclear ATXN3, Olig2, and MOBP. (B-D) No changes in total oligodendrocyte lineage cell counts (Olig2+) are seen, however immature oligodendrocytes (Mobp-/Olig2+ cells) are significantly increased, and mature oligodendrocytes (Mobp+) are significantly decreased in Q84/Q84 mice compared to wildtype. (E-F) ATXN3 nuclear accumulation does not change between Q84/Q84 and wildtype mice in immature oligodendrocytes but does significantly increase in mature oligodendrocytes in disease mice. Cell counts normalized to total DAPI-stained nuclei per field; ATXN3 nuclear accumulation normalized to wildtype images (n=3 images per mouse, n=4 mice per genotype). (G) Representative coronal CST transmission electron microscopy (TEM) images depict abnormal myelin wrapping in 16-week-old Q84/Q84 mice. Scale bar, 1μm. (H-J) Analysis of TEM images revealed an increase in g-ratio in Q84/Q84 mice, but no changes in axon caliber. Axon caliber and g-ratio were calculated using MyelTracer software^25^ (n=950 ± 150 axons per mouse, n=3 mice per genotype). Data reported as mean ± SEM. One-way ANOVA with Tukey’s multiple comparisons test or unpaired parametric t-tests, assuming equal SD, were performed (*p < 0.05, **p <0.01, ***p <0.001, ****p<0.0001, ns=not significant).

Transmission electron microscopy (TEM) images of the CST in homozygote Q84 SCA3 mice and wildtype littermates (**Fig 5G**) revealed ultrastructural abnormalities in the axonal myelination of disease mice at 16 weeks of age. Interestingly, there were no significant differences in axon caliber between genotypes (**Fig 5I**), however g-ratio analysis showed significant increases in Q84 CST compared to wildtype (**Fig 5H**). As g-ratio is calculated by the ratio of the inner axon diameter to the whole fiber diameter, higher g-ratios correspond to thinner myelin sheaths. When axon caliber and g-ratio are plotted together, thinner myelination around larger axons occurred significantly more in disease mice than in wildtype (**Fig. 5J**). Thus, we show that the reduction in mature oligodendrocytes has a functional consequence of thinner myelin sheaths in disease mice.

### Oligodendrocyte maturation impairment occurs early in SCA3 mouse disease pathogenesis

The above studies demonstrate impairment of oligodendrocyte maturation at the transcriptional, biochemical, and histological levels in vulnerable brain regions of SCA3 mice that translates to ultrastructural abnormalities in myelination, the main function of oligodendrocytes. To identify a timeframe during which dysfunction in oligodendrocyte differentiation begins, we transcriptionally assessed brainstem tissue at timepoints during which the highest rate of myelination is taking place^40^. We chose 4 weeks as it is the age of earliest onset of motor deficits (data not shown) and 3 weeks, because it is prior to any significant changes in motor activity. At 3 weeks of age, we see no differences in expression levels of oligodendrocyte maturation transcripts between wildtype and diseased mice (**Fig. 6A-D**). Interestingly, looking just one week later, the same trends seen at 8 weeks start to appear, with transcript levels of OPC marker *Smoc1* increasing (**Fig. 6A**) and expression of differentiating and mature oligodendrocyte markers (*Ugt8a*, *Plp1*, and *Mobp*) decreasing in Q84 mice compared to wildtype littermates (**Fig. 6B-D**). This result narrows the timeframe during which oligodendrocyte impairment begins to just seven days, giving further opportunity to investigate pathways affected early in disease and offering a timepoint after which therapeutic intervention could be less efficacious.

**Figure 6.**
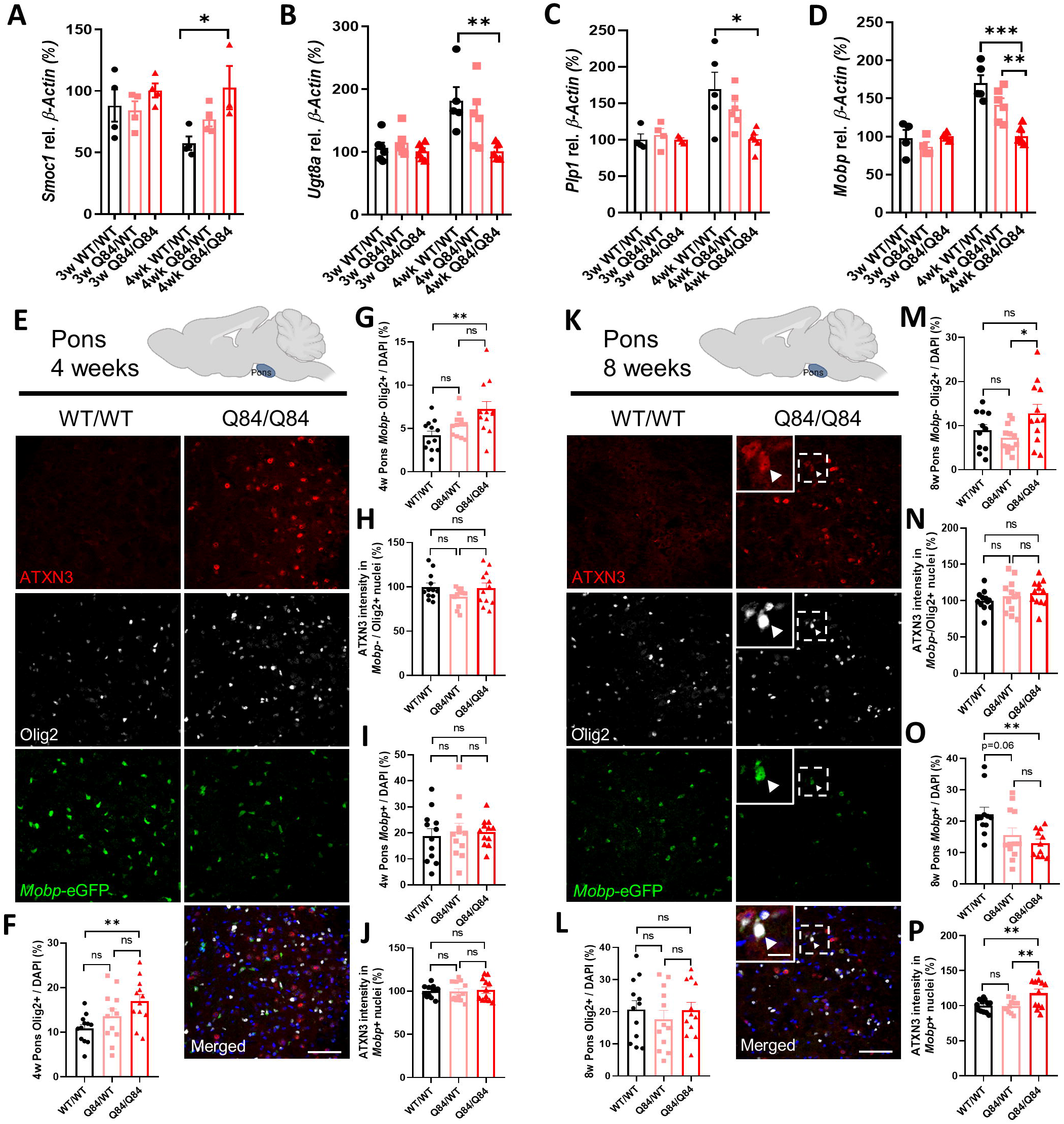
Oligodendrocyte transcript changes in the brainstem of SCA3 mice begin between 3 and 4 weeks of age, followed by protein changes between 4- and 8-weeks-old. (A-D) No changes in brainstem oligodendrocyte maturation transcripts are seen at 3 weeks, however by 4 weeks of age, there are significant increases in levels of OPC marker *Smoc1* and significant decreases in differentiating or mature oligodendrocyte markers *Ugt8a*, *Plp1,* and *Mobp*. (E, K) Representative sagittal immunofluorescent images of ATXN3 (red), Olig2 (white), and *Mobp*-eGFP (green) expression in 4-week-old and 8-week-old wildtype and Q84/Q84 mice in the body of the pons. Scale bar, 50 μm. Inset scale bar, 12.5 μm. White arrowheads define cells with colocalization of nuclear ATXN3, Olig2, and MOBP. (F-J) A significant increase in oligodendrocyte lineage cells, as well as immature oligodendrocytes, is seen at 4 weeks in the pons of Q84/Q84 mice, but there are no changes in mature oligodendrocyte cell counts or ATXN3 nuclear accumulation in immature or mature oligodendrocyte cells. (L-M) By 8 weeks of age, there are no significant differences in Olig2 positive cell counts or immature oligodendrocyte counts between Q84/Q84 and wildtype mice. (N) No changes in ATXN3 nuclear accumulation are observed in immature oligodendrocytes. (O-P) A significant reduction of *Mobp*-eGFP positive oligodendrocytes is seen at 8 weeks with increasing ATXN3 nuclear accumulation noted only in mature oligodendrocytes in disease mice. Cell counts normalized to total DAPI-stained nuclei per field; ATXN3 nuclear accumulation normalized to wildtype images (n=3 images per mouse, n= 4 mice per genotype). Data report as mean ± SEM. One-way ANOVA with Tukey’s multiple comparisons test was performed (*p < 0.05, **p <0.01, ***p <0.001, ns=not significant).

Histologically, mature oligodendrocyte transcriptional changes are not yet evident in 4-week-old pontine tissue, as defined by total Mobp+ cell counts (**Fig. 6E, I**). However, there is a significant increase in both total oligodendrocyte counts (Olig2+) and non-myelinating immature oligodendrocyte counts (Mobp-/Olig2+) in Q84 mice relative to wildtype mice (**Fig. 6F-G**), but with no differences in ATXN3 nuclear accumulation (**Fig. 6H, J**). By 8 weeks of age, total number of oligodendrocyte lineage cell counts is no longer significant across genotypes (**Fig. 6L**), but the changes in mature oligodendrocyte transcript levels at 4 weeks have translated into a significant reduction in mature oligodendrocyte counts in Q84 mice with a corresponding increase in nuclear ATXN3 (**Fig. 6K inset, O-P**). However, ATXN3 nuclear accumulation is still not present in immature oligodendrocytes at 8 weeks (**Fig. 6M-N**). These data indicate there may be a lag between transcript and protein dysregulation and that nuclear accumulation of ATXN3 may contribute to the impairment in oligodendrocyte maturation.

### Impairment of oligodendrocyte differentiation in SCA3 mice is cell autonomous

To determine if the deficit in oligodendrocyte maturation in Q84 mice is due to extrinsic factors, such as cell-to-cell signaling, or intrinsic factors, such as ATXN3 gain-of-function toxicity, we isolated and cultured OPCs from hemi- and homozygote Q84 mice alongside OPCs from WT littermates. Whole brains from post-natal day 5 (P5) to P7 mice were dissociated into a single cell suspension and immunopanned for the OPC-specific receptor PDGFRα using a magnetically conjugated antibody. Cells were run through a magnetic column, capturing labelled cells, which were then eluted and cultured in OPC proliferation media (+FGF2, +PDGFAA) for one day. Subsets of OPC cultured slides were either fixed prior to differentiation (DIV0), or switched to differentiation media (OPC media, −FGF2, −PDGFAA, +T3) for collection 3 days later (DIV3) (**Fig. 7A**).

**Figure 7.**
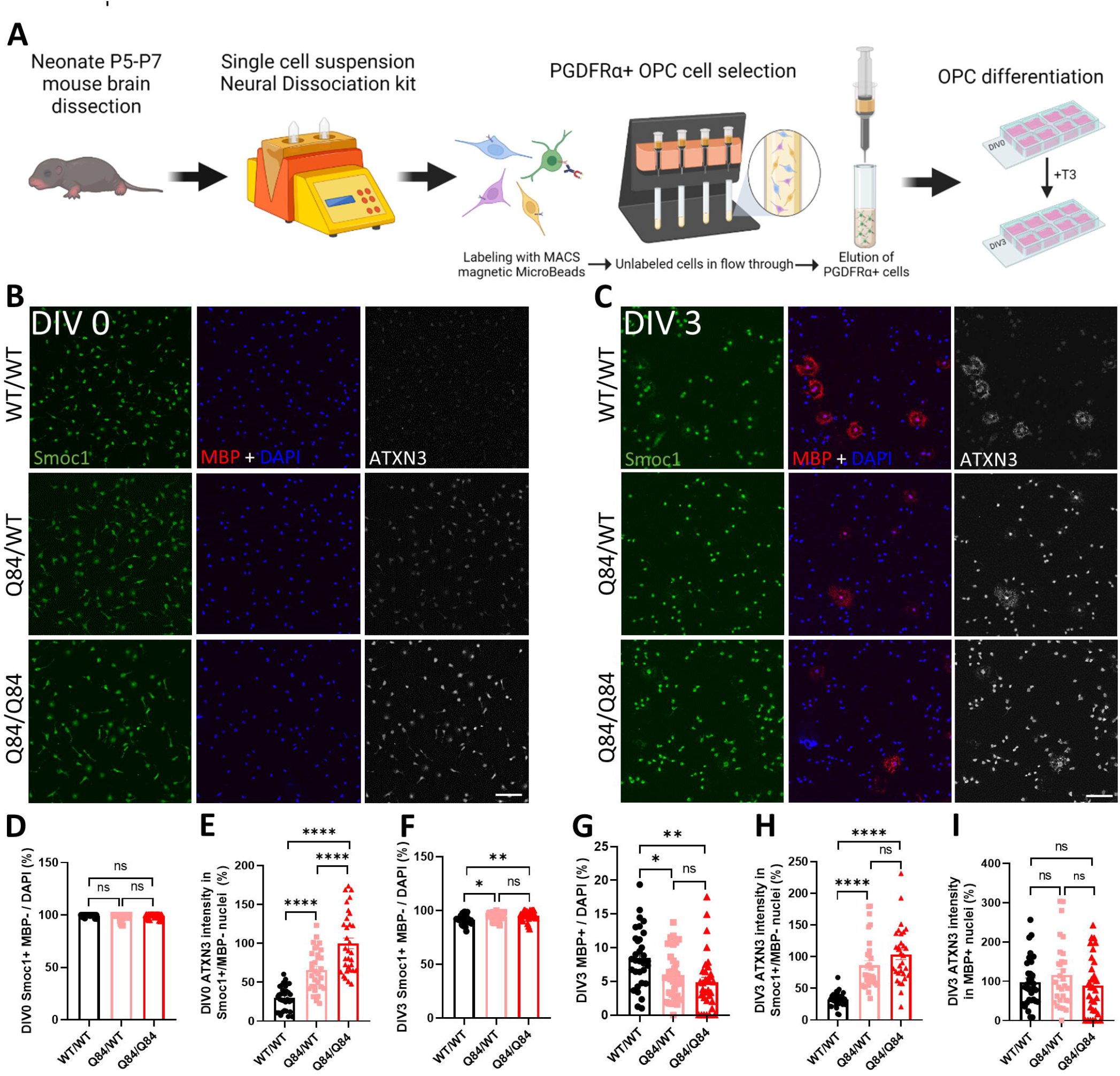
Cell-autonomous impairment of OPC differentiation to mature oligodendrocytes in cells cultured from SCA3 mice. (A) Schematic of OPC isolation and culture. (B-C) Representative immunofluorescent images of Smoc1 (green), MBP (red), ATXN3 (white), and DAPI (blue) expression in OPCs cultured from WT/WT, Q84/WT, and Q84/Q84 mice at DIV0 and DIV3. Scale bar, 100 μm. (D-E) Prior to differentiation, DIV0 OPC counts of Smoc1+/MBP− show no differences between genotypes, though cells cultured from disease mice display a dose dependent increase in ATXN3 nuclear accumulation. (F-G) After differentiating for 3 days in +T3 culture media (DIV3), there are significantly more immature (Smoc1+/MBP-) cells and fewer maturing (MBP+) cells in cultures from Q84/WT and Q84/Q84 mice. (H-I) ATXN3 accumulates significantly more in the nucleus of immature cells from disease mice, while maturing oligodendrocytes display no change of ATXN3 nuclear accumulation between genotypes. Cell counts normalized to total DAPI-stained nuclei per field; ATXN3 nuclear accumulation normalized to Q84/Q84 images (n=8 images per mouse, n=3-4 mice per genotype). Data report as mean ± SEM. One-way ANOVA with Tukey’s multiple comparisons test was performed (*p < 0.05, **p <0.01, ***p <0.001, ****p<0.0001, ns=not significant).

Oligodendrocytes at both timepoints were stained for the OPC marker SMOC1, the mature oligodendrocyte marker MBP, and ATXN3. (**Fig. 7B-C**). At DIV0, there were no differences in the number of OPCs (Smoc1+/MBP-), confirming that cells were plated at equivalent densities and at the same state of maturation (**Fig. 7D**). Interestingly, there is already a significant dose-dependent increase in the nuclear expression of ATXN3 in diseased OPCs at this time point (**Fig. 7E**). After 3 days of forced differentiation, there is an increase in Smoc1+/MBP− oligodendrocytes in the disease lines with a corresponding accumulation of nuclear ATXN3 in these immature cells (**Fig. 7F, H**). Conversely, there is a significant decrease in MBP+ mature oligodendrocytes by DIV3 with no changes in ATXN3 nuclear expression (**Fig. 7G, I**). This data demonstrates that when OPCs are removed from their normal environment and cultured independently in media to encourage differentiation, oligodendrocytes from diseased mice are unable to mature and those that remain immature are predisposed to a more severe nuclear deposition of ATXN3. The inability of primary OPCs from SCA3 mice to mature indicates that the impairment in oligodendrocyte differentiation is cell-autonomous and thus requires further investigation of the intrinsic mechanisms.

## DISCUSSION

The results here showing that oligodendrocyte dysfunction is an early and robust feature of SCA3 progression expand our understanding of disease pathogenesis in this polyglutamine disorder. Among SCA3 brainstem transcripts differentially expressed across all stages of disease, we found an enrichment in oligodendrocyte genes. These transcriptional changes are accompanied by evidence of nuclear accumulation of ATXN3 in oligodendrocyte lineage cells both *in vitro* and *in vivo*, corroborating a cell-autonomous effect. Using *ATXN3* KO mice, we verified that oligodendrocyte transcriptional changes in SCA3 mice were due to a toxic gain of function, not a loss of ATXN3 function. Through biochemical, histological, and ultrastructural analyses, we showed that the dysfunction of mature oligodendrocytes occurred in a region-specific pattern corresponding to brain areas known to be selectively vulnerable in SCA3, including the pons, DCN, and CST. Collectively our results indicate that, early in the SCA3 disease process, oligodendrocyte maturation is impaired in particularly vulnerable brain regions.

While some published evidence supports oligodendrocyte dysfunction in SCA3 disease, most of it comes from late-stage mouse models or post-mortem SCA3 patient tissue^9,16,17,41^, which fail to capture the context of oligodendrocyte dysfunction during the disease process. Here we define early impairments of oligodendrocyte maturation in an SCA3 mouse model that are cell autonomous and occur through a toxic gain of function mechanism. Our results place a spotlight on cells other than neurons early in SCA3 disease pathogenesis and strongly correlate with findings from another polyQ disease, HD. Recent studies in HD mouse models also revealed delayed oligodendrocyte differentiation^34,35^. Since *ATXN3* KO mice have no evidence of oligodendrocyte transcriptional or biochemical dysregulation^9^, this gain-of-function oligodendrocyte impairment may be more broadly implicated as a pathogenic mechanism in polyQ diseases. Future studies in these diseases should explore maturing oligodendrocyte populations for evidence of polyQ-related dysfunction.

There are also interesting similarities between SCA3 and HD as it pertains to cholesterol levels. Cholesterol is vital for many functions in the CNS, but in HD mouse models and human HD brain tissue, cholesterol levels are decreased due to a downregulation of genes involved in cholesterol biosynthesis^42^. Reduced cholesterol levels likely contribute to HD pathogenesis, based on the fact that mutant HTT-expressing embryonic rat striatal neurons supplemented with exogenous cholesterol are protected from cell death in a dose dependent manner^42^. Similar cholesterol dysfunction is found in SCA3 models. Through transcriptional profiling and subsequent application of IPA, Toonen *et al.* (2018) found cholesterol biosynthesis to be one of the top altered pathways in end-stage SCA3 mouse brainstem^17^. Moreover, a recent study in SCA3 mice found that overexpressing the rate-limiting enzyme for neuronally-synthesized cholesterol rescued SCA3 pathology and behavior^43^. Our IPA curated tox list indicated that genes involved in cholesterol biosynthesis may also be contributing significantly to oligodendrocyte dysregulation (**Fig 1E**), as cholesterol is necessary for oligodendrocyte-mediated axonal myelination in the developing brain and continued axon growth and synapse remodeling in the adult brain^44^.

In this study, we identified numerous dysregulated genes that relate to oligodendrocyte maturation. Proteolipid protein 1 (*Plp1)* in particular stands out as a key player, given its integral relationship to oligodendrocyte maturation and myelination. Since *Plp1* has not been associated with SCA3 disease pathogenesis before, we consider it to be among the genes that warrant further characterization in SCA3. *Plp1* encodes two major myelin proteins of the CNS, PLP and DM20 in oligodendrocytes. PLP is the most abundant integral membrane protein in myelin of the CNS, found almost exclusively in oligodendrocytes^45,46^. Both *Plp1* loss- and gain-of-function mutations can be detrimental to the brain. Loss-of-function mutations in *Plp1* elicit neuroinflammation, axonal degeneration, and neuronal loss^47^, whereas overexpression of *Plp1* can be damaging to the brain, as is found with incomplete penetrance in the neurodegenerative demyelinating Pelizaeus-Merzbacher disease (PMD)^48^. Exploring the effects of *Plp1* reduction in SCA3 disease could lead to a greater understanding of well-known SCA3 pathological features including neuroinflammation, axonal degeneration, and neuronal loss^1^.

Downregulation of *Plp1* in SCA3 may reflect dysregulation of the dual-regulatory TCF7L2 switch^29^. How this switch itself is regulated is not yet understood and would be interesting to resolve in the context of SCA3. TCF7L2 also functions in the peripheral nervous system (PNS) to control Schwann cell proliferation and regeneration^49^. Intriguingly, demyelination of the sciatic nerve in YACQ84 mice was one of the first described characteristics of this SCA3 mouse model in 2002^19^ and peripheral neuropathy is a common sign in SCA3 patients, due in part to demyelination^50^.

Research in SCA3 has focused on understanding neuronal dysfunction, but our findings combined with recent evidence from human and animal studies imply that glial cells have been a neglected contributor to SCA3 disease pathogenesis^1,9,17,43,51–53^. Among glial cell types in the mammalian brain, oligodendrocytes play integral roles in myelin-rich white matter structures that are vulnerable in SCA3^5,53^. Brain imaging now suggests that white matter changes in the brainstem are an early disease feature in human SCA3 disease^51–55^. In fact, other names for SCA3 have included spinopontine atrophy and spinocerebellar degeneration^56^, underscoring the central role of brainstem dysfunction in the disease. How these supporting glial cells contribute to SCA3 pathology is essentially unexplored. A better understanding of the mechanisms underlying oligodendrocyte impairments in SCA3 could shed further light on early steps in SCA3 pathogenesis and point to strategies for therapeutic intervention.

## ACKNOWLEDGMENTS

This work was supported in part by the National Ataxia Foundation SCA Young Investigator Award (H.S.M.), and grants from the NIH R01-NS122751 (H.S.M.), T32-NS007222-33 (H.S.M.), and U01-NS106670 (H.S.M. and H.L.P.). We acknowledge S. Meshinchi and D. Leroux in the Michigan Medicine Microscopy Core for TEM training and processing of TEM samples. Figure schematics created using BioRender.com.

